# Symptomatic human norovirus infection in zebrafish embryos uncovers neural infection and extracellular vesicle-mediated transmission dynamics

**DOI:** 10.64898/2026.06.22.733919

**Authors:** Malcolm Turk Hsern Tan, Huixin Duan, Zejia Lin, Jillinda Yi Ling Toh, Hong Bai, Kun Qu, Dan Li

## Abstract

Human norovirus (hNoV) is the leading global cause of acute gastroenteritis, imposing a substantial health and economic burden worldwide. Progress in understanding hNoV pathogenesis has been hindered by the lack of tractable small-animal models that recapitulate symptomatic infection. Although zebrafish larvae support hNoV replication, infection remains asymptomatic, limiting their utility for studying disease mechanisms and host-pathogen interactions. In this study, we report that the zebrafish embryo infection model, in which microinjection of hNoV at the early cell stage, resulted in robust systemic viral replication accompanied by overt pathological manifestations, including pericardial and renal edema, yolk and cranial opacity, and mortality by 3 days post-infection. Disease severity displayed marked individual variability and correlated closely with viral burden. Integrated multi-omics analyses, including bulk transcriptomics, untargeted metabolomics, and single-cell RNA sequencing, demonstrated that embryonic infection elicits a stronger and more coordinated antiviral response than larval-stage infection, while enabling widespread viral dissemination across diverse cell lineages. Approximately two-thirds of infected cells were derived from the nervous system or neural crest lineages, providing a potential mechanistic basis for the neurological complications occasionally reported in hNoV-infected patients. Furthermore, we identified a developmental stage-dependent role for extracellular vesicle (EV)-associated hNoV transmission: free virions mediated more efficient infection and higher symptomatic incidence in immunologically immature embryos, whereas EV-associated virions exhibited enhanced infectivity in more immunocompetent larvae. Together, these findings establish the zebrafish embryo as a versatile and accessible *in vivo* platform for studying symptomatic hNoV infection, reveal host maturity-dependent viral transmission strategies, and provide new opportunities for mechanistic studies and high-throughput evaluation of antiviral and vaccine candidates.

**Author summary:** Human norovirus is the leading cause of stomach flu worldwide but studying it has been difficult because the lack of a simple, small-animal model that actually gets sick from the virus. While older zebrafish larvae can harbor the virus, they do not show symptoms. In this study, we successfully created a new model by injecting human norovirus into zebrafish embryos at their early cell stage. Unlike the older larvae, these embryos developed clear symptoms, including fluid buildup around the heart and kidneys, tissue cloudiness, and death within three days. Using advanced genetic and metabolic tracking, we discovered that the virus spreads widely throughout the embryo’s body, particularly targeting cells in the nervous system. This link might explain why human patients occasionally suffer from neurological symptoms. Additionally, our study revealed that the virus changes its transmission strategy based on the animal’s age: it travels freely to infect vulnerable embryos but hides inside lipid vesicles to infect older, more immune-developed larvae. Ultimately, these findings provide a practical, efficient animal model to better understand norovirus sickness and rapidly test new vaccines and treatments.

## Introduction

Human norovirus (hNoV) is recognized as the leading cause of acute gastroenteritis worldwide and represents a substantial global public health burden. It is estimated that hNoV causes approximately 685 million cases of acute gastroenteritis annually, including ∼200 million infections among children under five years of age, resulting in roughly 200,000 deaths globally each year, with the majority occurring in low-income countries and vulnerable populations such as young children and the elderly (WHO, 2015). The virus is responsible for about one in five cases of acute gastroenteritis worldwide, highlighting its dominant role among enteric pathogens (Hashemi et al., 2023). Beyond the clinical burden, hNoV also imposes a major economic impact, with global costs estimated at around US$60 billion annually due to healthcare expenditures and productivity losses (Saleem et al., 2025). In addition, hNoV is the leading cause of foodborne illness in many developed countries, accounting for more than half of foodborne disease outbreaks in the United States alone (CDC, 2026).

HNoV infections are typically acute and self-limiting, characterized by the sudden onset of nausea, vomiting, diarrhea, and abdominal pain. Symptoms usually appear within 12-48 hours after exposure and generally resolve within 1-3 days in otherwise healthy individuals without the need for specific medical treatment. Despite the generally mild clinical course, hNoV infections can pose a significant health risk to vulnerable populations, including elderly individuals, young children, and immunocompromised patients. In these groups, the illness may be prolonged and more severe, which may require hospitalization. Epidemiological studies have shown that older adults with underlying comorbidities are particularly susceptible to severe outcomes, with increased risks of hospitalization and short-term mortality following laboratory-confirmed hNoV infection (Cates et al., 2023).

Although hNoV was first identified in the 1970s (Kapikian et al., 1972), its pathogenesis has long remained poorly understood due to the historical lack of a robust permissive cell culture system and suitable animal models for studying viral infection. This limitation significantly hindered mechanistic investigations of viral replication, host interaction, and disease progression for decades. In the absence of reliable experimental systems, researchers have often relied on surrogate viruses that can be propagated in cell culture to approximate the behavior of hNoV. Commonly used surrogates include feline calicivirus (FCV), murine norovirus (MNV), and Tulane virus (TV). FCV was historically the earliest surrogate used because it belongs to the same family (*Caliciviridae*) and can be readily cultured (Radford et al., 2007), while MNV later became widely adopted after its discovery in 2003 due to its genetic and structural similarities to hNoV and its ability to replicate in cell culture and small animal models (Wobus et al., 2006). More recently, Tulane virus, a *Recovirus* isolated from rhesus macaques, has been proposed as another surrogate because it recognizes histo-blood group antigens similarly to hNoV (Farkas, 2015). Nevertheless, these surrogate viruses only partially mimic the biology of hNoV and often fail to reproduce key features of human infection, particularly the clinical symptoms observed in patients. Attempts to establish animal models for hNoV using species such as chimpanzees (Bok et al., 2011), gnotobiotic pigs (Cheetham et al., 2006), and gnotobiotic calves (Souza et al., 2008) have resulted in only limited viral replication and have rarely reproduced the characteristic symptoms of hNoV caused gastroenteritis.

Only with the advent of the human intestinal enteroid (HIE) culture system have researchers, for the first time, been able to successfully cultivate hNoV in an *in vitro* laboratory setting. This breakthrough, first reported in 2016, demonstrated that stem cell derived intestinal epithelial cultures can support the replication of multiple hNoV genotypes, overcoming a decades-long barrier in hNoV research (Ettayebi et al., 2016). The HIE model closely recapitulates the cellular composition and physiology of the human intestinal epithelium and has since become a powerful platform for investigating viral replication, host pathogen interactions, antiviral responses, and virus inactivation strategies (Costantini et al., 2025; Murakami et al., 2020; Prasad et al., 2025).

Shortly thereafter, zebrafish (*Danio rerio*) larvae were identified as a permissive *in vivo* model for hNoV infection, providing a simple and tractable vertebrate host system for studying viral replication and antiviral interventions (Van Dycke et al., 2019). Studies have shown that multiple genotypes of hNoV can replicate efficiently in zebrafish larvae following microinjection, with viral replication detectable for several days post-infection. This model offers several advantages, including optical transparency, ease of handling, and suitability for high-throughput infection studies (Van Dycke et al., 2021). Further investigations conducted by our research group have revealed that hNoV infectivity in zebrafish is developmentally stage-dependent, with embryos supporting higher levels of viral replication than larvae. Experimental comparisons have shown that virus replication in zebrafish embryos can be detected earlier, producing higher viral loads and enabling continuous viral passaging (Tan et al., 2023). While recent advances have utilized the zebrafish embryo as a platform for reverse genetics (Kotaki et al. 2025), the underlying mechanisms of tissue tropism and the developmental factors governing symptomatic infection remain poorly understood.

In this study, we present a multiscale characterization of hNoV pathogenesis in zebrafish embryos by integrating transcriptomic, metabolomic, and single-cell analyses. This comprehensive approach was used to evaluate the extent to which the zebrafish embryo model recapitulates key features of human infection, with particular emphasis on symptomatic infection and the observed inter-individual variability in disease outcomes. In addition, we identified a pronounced production of extracellular vesicles during hNoV infection in zebrafish embryos and further investigated their potential role in hNoV infection and host-virus interactions.

## Results

### Transcriptomic analysis and untargeted metabolomics

Compared with the infection achieved by injecting hNoV into zebrafish larvae at 3 dpf (Figure 1A), a more pronounced infection was observed when hNoV GII.4[P16] (∼100 genome copies per embryo) was delivered to zebrafish embryos at 0 day post-fertilization (dpf) (Figure 1B). Bulk RNA-sequencing analysis was performed on hNoV-infected larvae at 3 days post-injection (dpi; injection conducted at 0 dpf; 10 larvae randomly pooled per sample), corresponding to the peak viral load as reported previously (Tan et al., 2023). A total of 494 genes were differentially expressed, including 381 upregulated and 113 downregulated genes (Supplementary 1). On the contrary, when the same experimental design (10 larvae randomly pooled per sample) was applied to zebrafish larvae infected at 3 dpf and analyzed at 2 dpi, when viral load peaks under this infection regime (Tan et al., 2023), only 45 genes were significantly upregulated compared with the mock-injected controls (Tan et al., 2022). This marked difference in transcriptional responses suggests that infection at the embryonic stage triggers a substantially stronger host response than infection initiated at the larval stage.

**Figure 1.**
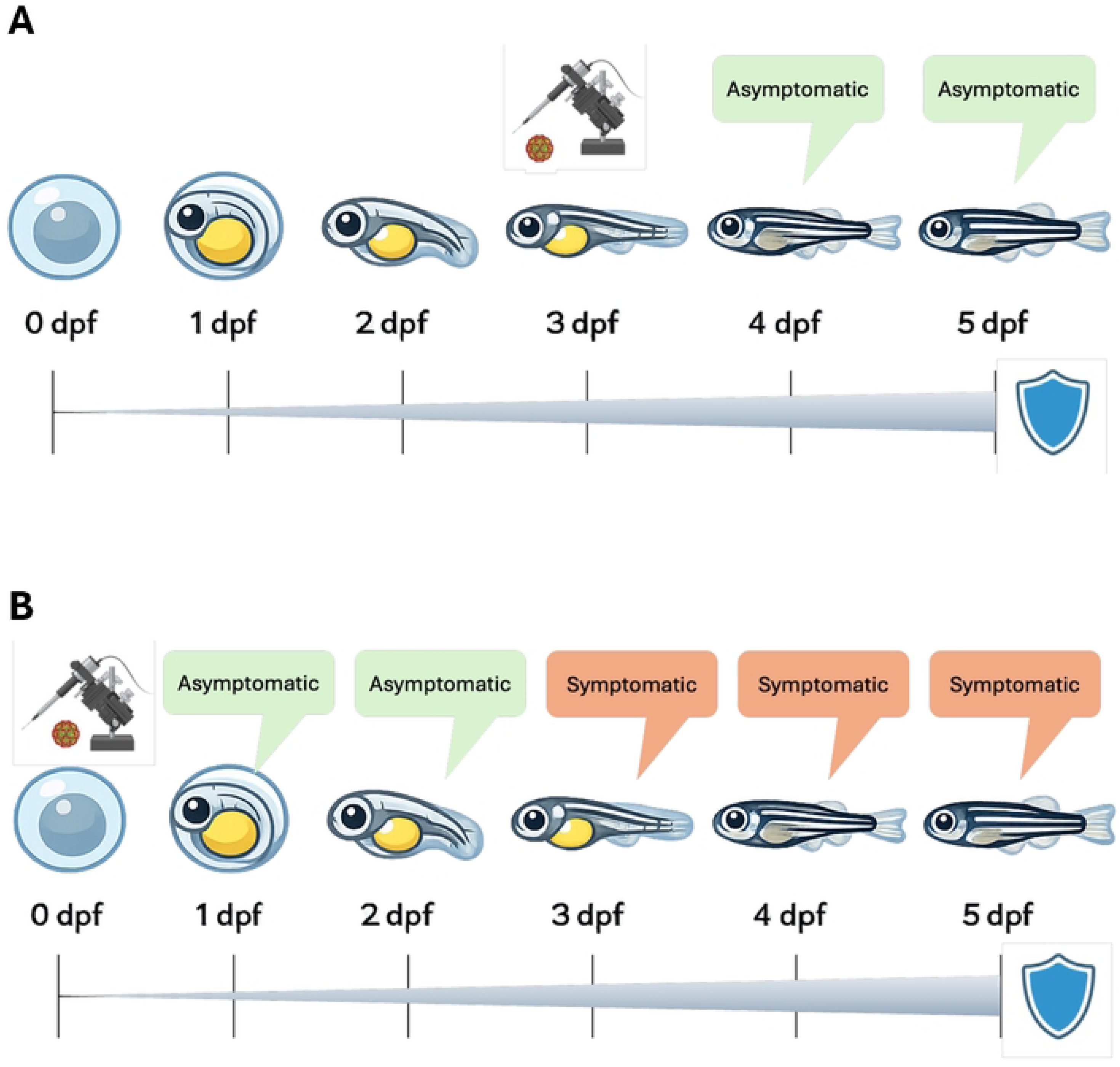
Experimental timeline and pathological outcome following hNoV exposure in zebrafish embryos and larvae. **(A)** Schematic representation of hNoV exposure during the larval stage. At 3 dpf, larvae are exposed to virus. Following exposure, individuals remain asymptomatic at 4 dpf and 5 dpf. **(B)** Schematic representation of hNoV exposure during the embryonic stage. Embryos are exposed to virus at 4 hpf. Individuals remain asymptomatic at 1-2 dpf, but begin to display pathological phenotypes from 3 dpf onward.

Analysis of the differentially expressed genes (DEGs) using ShinyGO showed predominant enrichment of KEGG pathways related to immune responses (Figure 2A). Notably, the RIG-I-like receptor signalling and Toll-like receptor signalling pathways were among the top enriched pathways, indicating an upregulation of the pathogen recognition system in response to viral infection (Kawai & Akira, 2010; Yoneyama & Fujita, 2008). The phagosome pathway, which mediates pathogen clearance, was similarly enriched in infected larvae (Flannagan et al., 2011). Additionally, the enrichment of cytokine-cytokine receptor interaction and FoxO signalling pathways suggests downstream modulation of inflammatory responses and cellular stress adaptation (Kim et al., 2022; O’shea et al., 2019). Interestingly, 20 ribosomal genes were downregulated (Supplementary 1), pointing toward a reduction in overall protein synthesis that may serve as a host defence strategy to limit viral replication while preserving resources for immune activation and stress responses (Walsh & Mohr, 2011). Together, these findings indicate that infected larvae mounted a coordinated antiviral response, simultaneously activating immune defences while modulating ribosomal activity and protein synthesis to prioritize host survival.

**Figure 2.**
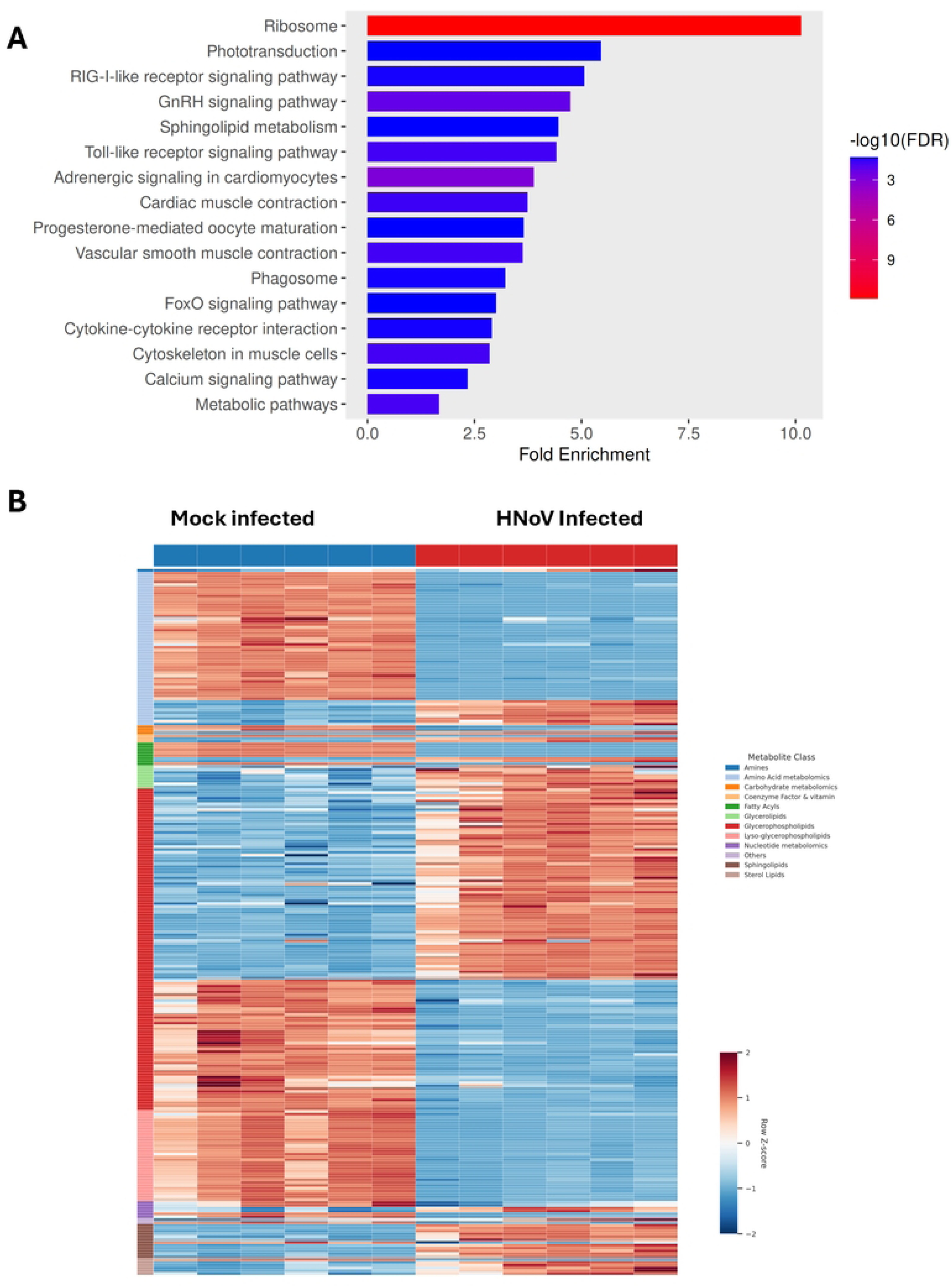
Human norovirus infection induces coordinated transcriptomic and metabolomic remodelling. **(A)** KEGG pathway enrichment analysis of differentially expressed genes between mock and hNoV-injected groups. **(B)** Heatmap of differentially abundant metabolites across biological replicates from mock and hNoV injected groups.

The transcriptomic data also revealed enrichment of metabolic pathways, which prompted subsequent untargeted metabolomic profiling to explore whether these transcriptional changes were reflected at the metabolite level. Compared with the mock-injected group, hNoV-infected larvae at 3 dpf (injection conducted at 0 dpf; 10 larvae randomly pooled per sample) exhibited widespread shifts in metabolite abundance, mainly among lipids, amino acids, and carbohydrate-related metabolites, suggesting a substantial metabolic remodelling in response to infection (Figure 2B). Alternations in lipids comprised the largest portion of the metabolite changes, highlighting the extensive influence of hNoV infection on lipid metabolism. Sphingolipids and glycerophospholipids, which represented the majority of altered lipids, are critical for membrane structure and signalling, supporting the notion that hNoV infection may impact these processes (Heaton & Randall, 2011). Sphingolipids, in particular, showed concordant changes across both metabolomic and transcriptomic pathway enrichment analyses, highlighting their potential functional involvement during hNoV infection.

### Embryos injected with hNoV exhibit pathological symptoms upon reaching the larval stage with individual variances

Being clearly different with the zebrafish larvae model which results in only asymptomatic virus replication (Figure 1A), we observed that hNoV embryonic-stage infection typically resulted in the development of pathological symptoms, with onset occurring as early as 3 dpi (Figure 1B). Phenotypically, affected larvae exhibited edema localized to the pericardial region, renal region, or both, as well as opacity appearing on the yolk, head, or both (Figure 3A, II-III). In more severe cases, larvae were found dead, exhibiting extensive tissue degradation or complete loss of structural integrity (Figure 3A, IV). We next found that the pathological symptoms correlated closely with viral loads (Figure 3B). Symptomatic individuals consistently harboured higher viral loads (7.2 ± 0.7 log genomic copies/larva), whereas asymptomatic individuals carried significantly lower (2.6 ± 2.2 log genomic copies/larva, *P* < 0.05) viral loads. In most asymptomatic individuals, hNoV levels remained comparable to the initial exposure level or below the detection limit (< 1 log genome copies/larva) and showed substantially greater inter-individual variances. Nonetheless, high viral loads were occasionally observed among the asymptomatic fish, particularly following high-dose exposure on 0 dpf (Figure 3B). These phenotypic manifestations offer a simple visual proxy for viral burden, linking observable pathology to viral infectivity.

**Figure 3.**
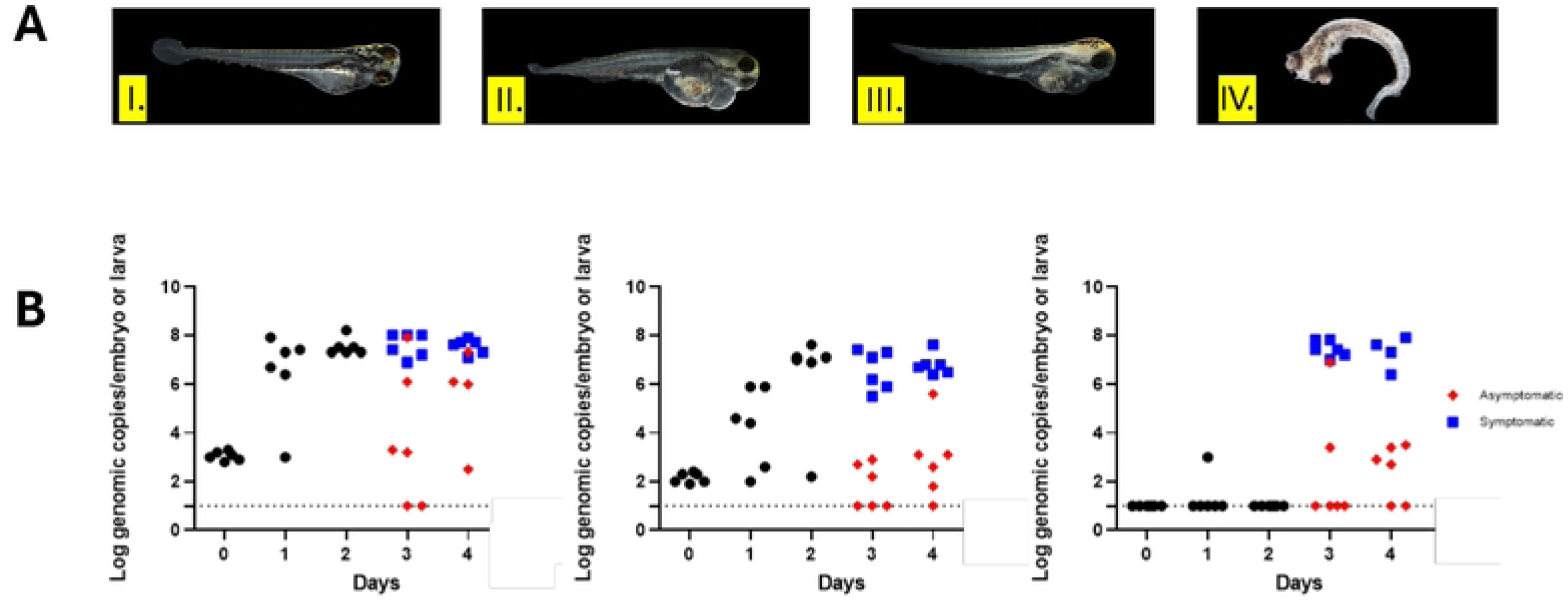
**(A)** Representative pathological phenotypes observed following infection with hNoV: (1) asymptomatic larva showing no observable clinical sign, (2,3) symptomatic larva exhibiting (a) edema localized to pericardial region, renal region, or both as well as (b) opacity appearing on the yolk, head, or (c) both, (3) dead larva displaying extensive tissue degradation. **(B)** Human norovirus genomic copies (GC) quantified in individual zebrafish embryo/larva post-infection following microinjection with: (1) 1,000 GC/embryo, (2) 100 GC/embryo, and (3) 10 GC/embryo.

### Single-cell RNA sequencing (scRNA-seq) and clustering

scRNA-seq was employed as an attempt to identify individual cells that may serve as sites for hNoV replication in zebrafish and to obtain a more comprehensive view of differential gene expression. Based on the established evidence that pathologically symptomatic individuals are known to harbour higher viral loads, scRNA-seq was performed exclusively by pooling symptomatic individuals. This targeted approach was designed to avoid pooling individuals with low viral loads, which could dilute the signal and reduce the likelihood of identifying virus-infected cell populations. In the mock-injected control group, 12,832 cells met the inclusion criteria, with a mean of 32,065 reads per cell and a median of 1,198 genes per cell. On the other hand, the hNoV-infected group had 26,874 qualifying cells, with a mean of 15,172 reads per cell and a median of 824 genes per cell. A summary of the scRNA-seq results can be found in Supplementary 2. Using the expression data from the top 16 most differentially expressed genes previously annotated by Farnsworth et al. (2019), 25 distinct cell clusters were identified. These cluster are visualized in Figure 4A as a uniform manifold approximation and projection (UMAP) plot. The genes used for assignment of each cluster are shown in Supplementary 3.

**Figure 4.**
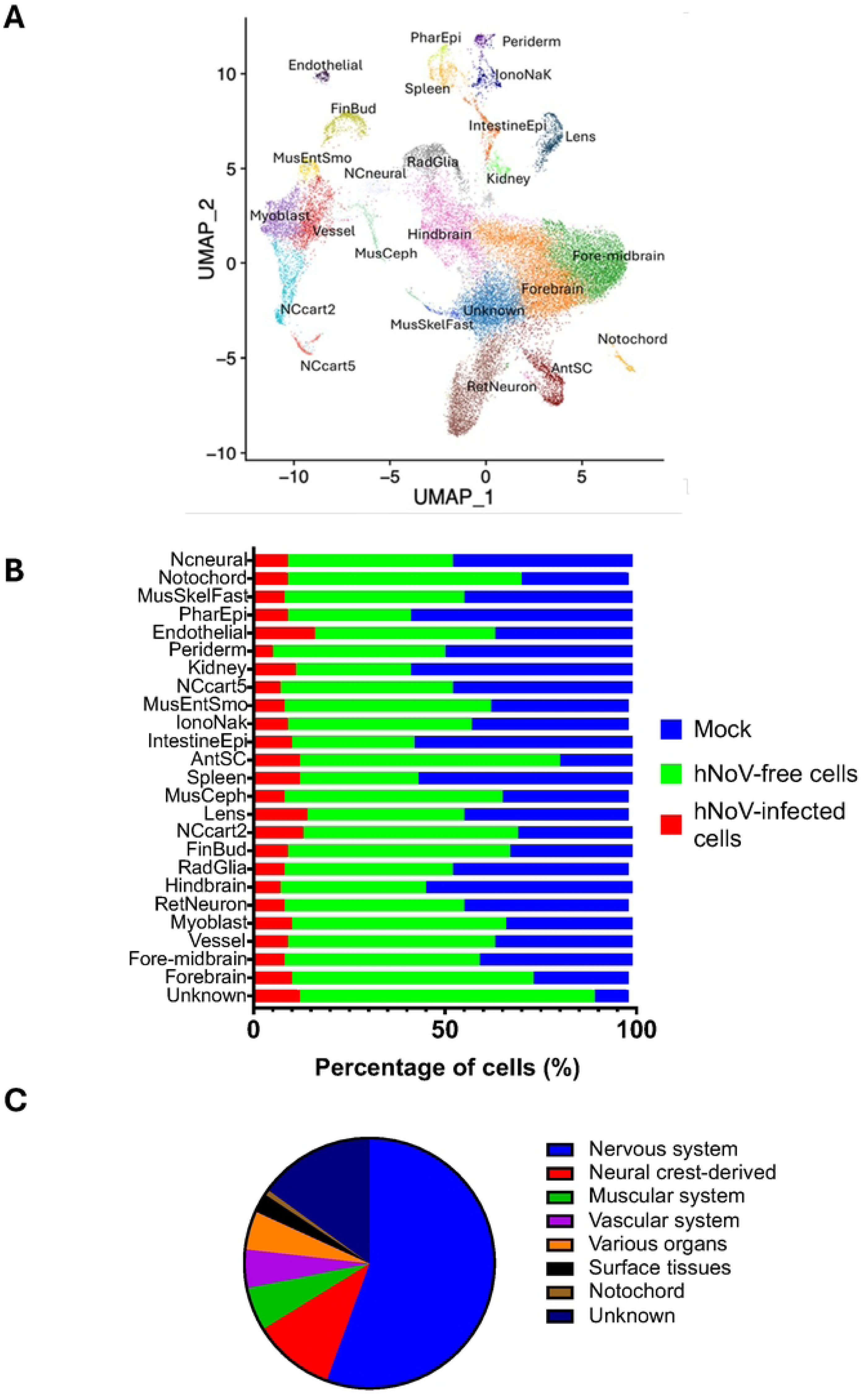
Single-cell transcriptomic analysis of host response to hNoV infection. **(A)** Uniform Manifold Approximation and Projection (UMAP) visualisation data showing clustering and annotation of cell types. **(B)** Proportional distribution of annotated cells stratified by infection status. **(C)** Distribution of viral RNA across the distinct cell categories, showing which major cell groups harboring the virus.

The scRNA-seq data were screened for hNoV expression to determine whether the virus localised to specific clusters. A total of 3,169 (15.5%) out of 20,508 cells were found to express the viral RNA, with the expression distributed across all the identified the clusters (Figure 4B). The widespread presence and heterogenous distribution of hNoV transcripts suggests that the infection was systemic in symptomatic individuals.

The clusters in Figure 4A and 4B were further simplified into distinct categories: the nervous system-associated cells (including the forebrain, fore-midbrain, hindbrain, retinal neurons, radial glial cells, and anterior spinal cord), neural crest-derived cells (such as neural crest cartilage, cephalic muscle, lens, pharyngeal epithelium, and neural crest neural), the muscular system (myoblasts, fast skeletal muscle, and enteric smooth muscle), the vascular system (vessels and endothelial cells), various organs (kidney, spleen, intestinal epithelium, and ionocytes), surface tissues (periderm and finbud), the notochord, and an unidentified cluster (Figure 4C). Majority of hNoV infected cells were of neural origin, including both cells from the nervous system-associated cells (55.6%) and neural crest-derived populations (10.6%) (Figure 4C). The remaining of the categories represented 19.8% of the total. The unidentified cluster accounted for 15% of the remaining viral transcripts. The cells within this cluster may have lost their original lineage markers and could also be neuron-related, as evidenced by their close spatial clustering and low expression of some brain specific markers (data not shown). Consistent with the bulk RNA-seq results, genes associated with the innate immune response were upregulated across multiple clusters. However, high expression of hNoV RNA and interferon-stimulated genes (ISGs) did not necessarily occur within the same clusters of individual cells. Specifically, as shown in Figure 5, the blue framed clusters represent cells expressing the highest levels of ISGs, including *isg15, rsad2, ifi45,* and *ifi46*, whereas these clusters did not correspond to those with the highest levels of hNoV RNA. Similarly, the red framed clusters represent cells expressing the highest levels of hNoV RNA, whereas these clusters did not correspond to those with the highest levels of ISGs.

**Figure 5.**
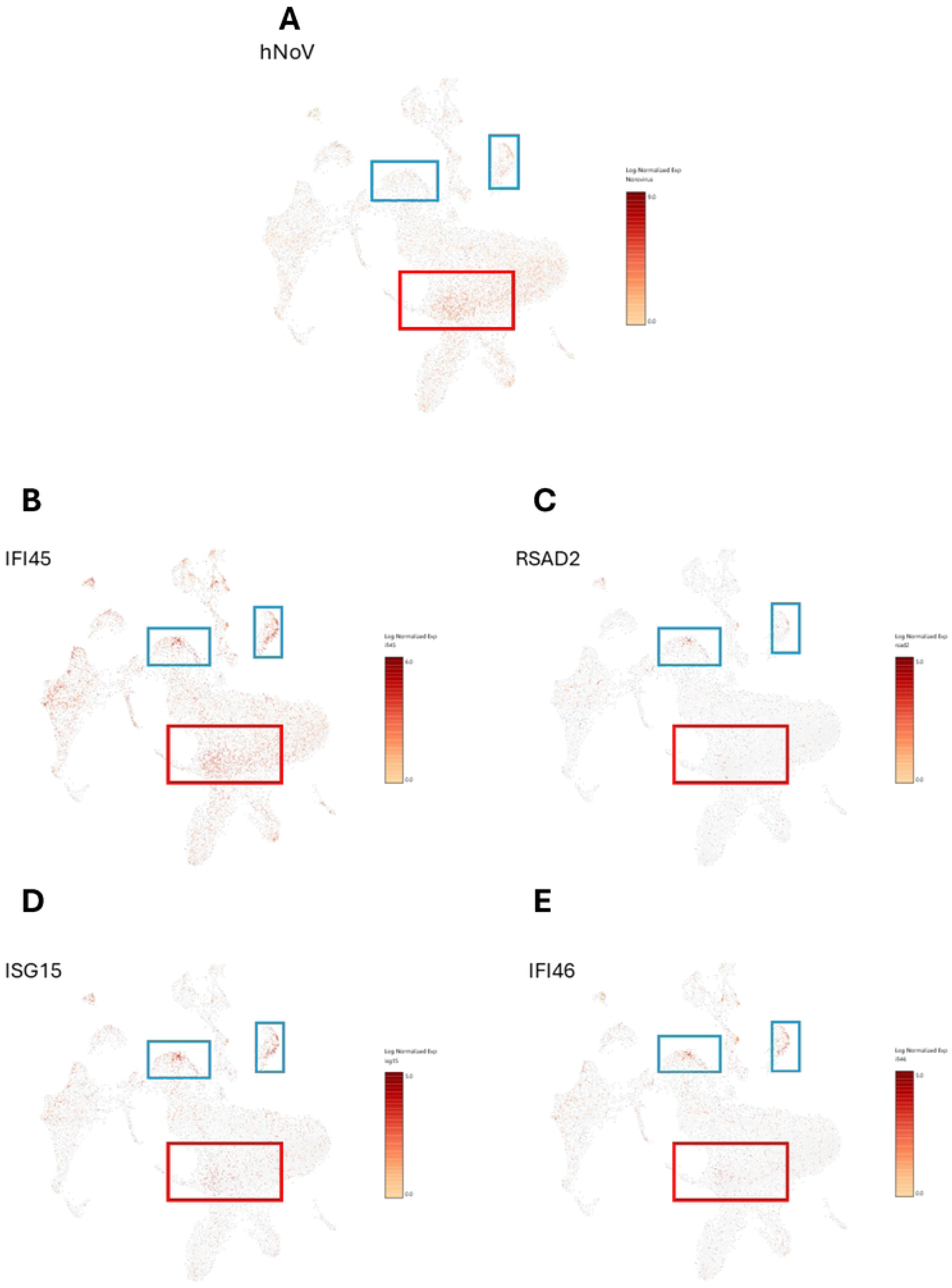
**(A)** Distribution of hNoV viral RNA reads across individual cells. Feature plots showing the expression of interferon-stimulated genes **(B)** ifi45, **(C)** RSAD2, **(D)** ISG15 and **(E)** ifi46 in hNoV infected larvae. Cells expressing the genes are highlighted in red, indicating activation of immune response in specific cell populations following viral infection.

### Altered barrier integrity in hNoV infected larvae

As hNoV transcripts were previously observed to be widely distributed throughout infected larvae, we investigated whether the viral infection was associated with altered tissue permeability. To address this, 10 kda dextran conjugated with Alexa Fluor™ 647 (dextran-AF647) was used as a fluorescent tracer; its intermediate molecular weight allows it to remain largely excluded from intact biological barriers while becoming detectable in cases of increased permeability (Meirelles et al., 2025). Figure 6 shows representative brightfield and Texas Red fluorescence images of mock-injected and hNoV-infected larvae, with or without dextran-AF647. Larvae that did not receive dextran-AF647 showed no detectable background fluorescence in either group (Figure 6A and 6B for Texas Red). On the other hand, dextran-AF647 injected groups exhibited strong fluorescence signal. In mock-injected larvae, fluorescence was largely restricted to the yolk sac (Figure 6C for Texas Red), whereas hNoV-infected larvae displayed a widespread distribution of fluorescence throughout the body (Figure 6D for Texas Red). These results suggest that hNoV infection is associated with increased in tissue permeability and/or disruption of barrier integrity, allowing dextran-AF647 to spread beyond its expected confinement.

**Figure 6.**
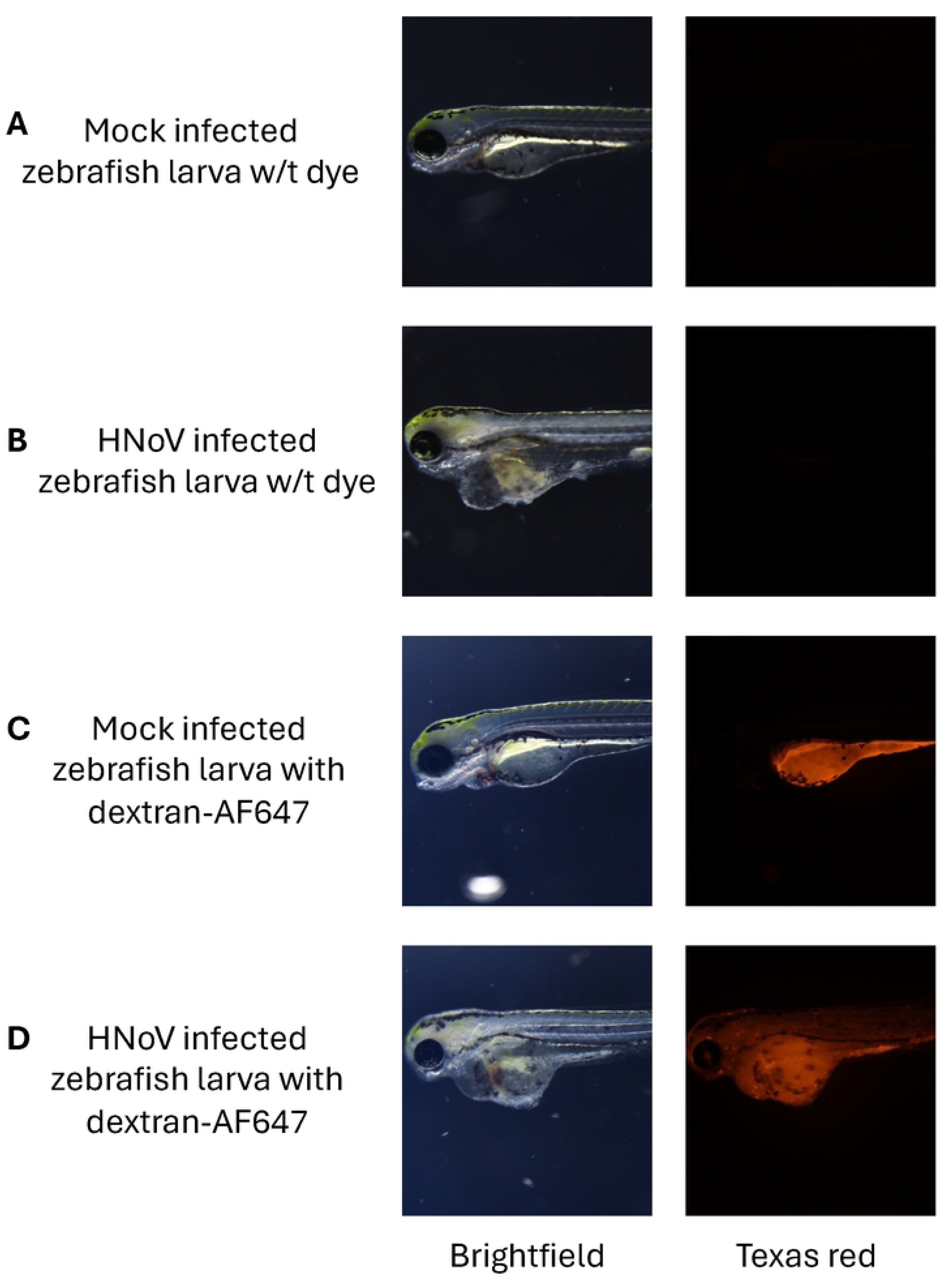
Distribution of dextran-AF647 in mock-injected and hNoV-infected zebrafish larvae. Representative brightfield (left) and Texas Red (right) fluorescence images of zebrafish larvae following mock infection or hNoV infection. Larvae without dextran-AF647 **(A)** and **(B)** showed no background fluorescence in Texas Red channel. In contrast, larvae injected with dextran-AF647 exhibited strong red fluorescence signal; **(C)** mock-injected larvae displayed localised fluorescence primarily in the yolk sac region, whereas **(D)** hNoV-infected larvae showed more widespread and intense across the body.

### Role of extracellular vesicles (EVs) in hNoV infection in zebrafish

The long-held assumption that hNoV exists solely as free viral particles was challenged by Santiana et al. (2018), who demonstrated that the virus could occur in both free and vesicle-bound forms in human stool samples. Here, for the first time, we show that hNoVs can also exist in vesicle-bound form in zebrafish. This conclusion was first supported by cryo-electron microscopy images of hNoV produced in zebrafish embryos/larvae, which revealed membranous vesicle-like structures of approximately ∼50 nm in diameter alongside free viral particles of ∼30 nm in diameter (Figure 7A). Next, we incubated the hNoVs harvested from zebrafish embryos/larvae with TIM4-coupled beads, which was identified as a phosphatidylserine receptor and commonly used to recover EVs (Miyanishi et al., 2007; Santiana et al., 2018). The hNoVs in EVs recovered by the TIM4-coupled beads and the non-bound free viral particles were enumerated with the use of reverse transcription-quantitative PCR (RT-qPCR) respectively. As shown in Figure 7B, the vesicle-bound viruses accounted for approximately 26% of the total viral population. Reductions in vesicle-bound viruses were observed after 5 and 10 freeze-thaw cycles, which was expected to compromise vesicle integrity and consequently release more free viral particles (Figure 7B).

**Figure 7.**
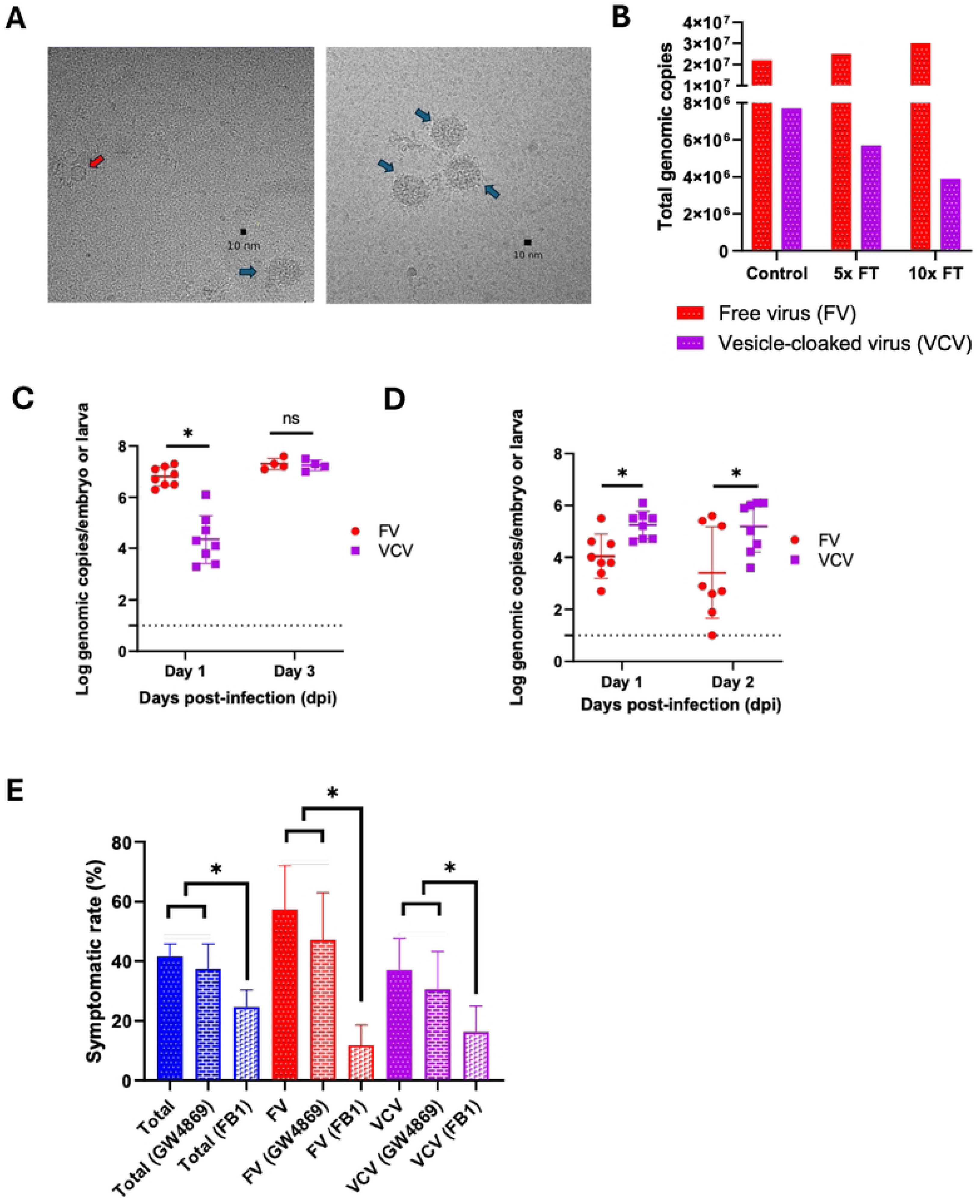
**(A)** Representative transmission electron micrograph showing free norovirus particle (red arrow) and vesicle-bound norovirus (blue arrow). Scale bars represent 10nm. **(B)** Quantification of free virus particles and vesicle cloaked norovirus following TIM4 bead treatment. Samples were quantified under control condition and after 5x and 10x free-thaw cycles. Red bars represent free virus particles, while purple bars represent vesicle-cloaked virus. Total genomic copies detected in individual embryo/larva infected with free virus (red) and vesicle-cloaked virus (purple). Infection was performed at two developmental stages: **(C)** embryo stage, **(D)** larval stage (3 days post-fertilization). **(E)** Effect of ceramide inhibitor treatment on the percentage of symptomatic individuals following hNoV exposure. Blue bars represent infection with unfractionated virus (total), red bars represent infection with free virus particles and purple represent infection with vesicle-cloaked virus.

Moving forward, it was interesting to find that the role of EVs in hNoV infection in zebrafish was stage dependent. Each embryo or larva was injected with ∼100 genome copies of either vesicle-bound or free hNoVs. For zebrafish embryos injected with hNoV on 0 dpf, on 1 dpi, the viral loads in embryos injected with free viruses were significantly higher than those injected with vesicle-bound viruses (*P* < 0.05; Figure 7C). On 3 dpi, the viral loads were equally high for both vesicle-bound and free viruses in pathologically symptomatic larvae (*P* > 0.05; Figure 7C). However, as demonstrated above, the virus loads in symptomatic larvae were significantly higher than asymptomatic larvae on 3 dpi (*P* < 0.05; Figure 3B). In this test, it was noticed that the symptomatic rate of zebrafish larvae on 3 dpi was significantly higher after injection of free viruses than vesicle-bound virus on 0 dpf (*P* < 0.05; Figure 7E). In contrast, for zebrafish larvae injected with hNoV on 3 dpf, free viruses displayed less efficient infectivity compared to vesicle-cloaked viruses on both 1 dpi and 2 dpi (*P* < 0.05; Figure 7D).

GW4869 is a non-competitive inhibitor of neutral sphingomyelinase that is widely used in research to block exosome biogenesis and release (Murakami et al., 2020). In this study, treatment with GW4869 did not result in significant differences in symptomatic rates in zebrafish larvae at 3 dpi following injection at 0 dpf with free viruses, vesicle-bound viruses, or their mixture (*P* > 0.05; Figure 7E). In contrast, injection of fumonisin B1 (FB1), an inhibitor of ceramide synthases as the key enzymes in the *de novo* ceramide biosynthesis pathway, led to significant reductions in symptomatic rates in zebrafish larvae at 3 dpi after injection of free viruses, vesicle-bound viruses, or their mixture at 0 dpf (*P* < 0.05; Figure 7E).

## Discussion

One of the key challenges of the zebrafish larval-stage infection model is that hNoV replication does not induce observable signs of distress or disease, making it difficult to visually distinguish infected from uninfected individuals. Consequently, viral load quantification has relied primarily on RT-qPCR, a method that is both costly and time-consuming. In practice, larvae are typically pooled in groups of ten to obtain an average viral load. Because pooling is performed randomly, each pool likely contains a heterogeneous mixture of infected and uninfected larvae, which may dilute the measured signal and limit resolution at the individual level. In contrast, embryonic-stage infection, as demonstrated in this study, exhibited clear and discernible signs of infection. Visible symptoms were observed in larvae as early as 3 dpi following hNoV infection. The strong correlation between viral burden and phenotypic changes opens new avenues for establishing a straightforward approach to assess viral infectivity based on symptom prevalence.

Timing is critical for generating the unique symptomatic hNoV infection in zebrafish embryos described in this study. Early embryonic development offers a highly permissive environment for hNoV replication in zebrafish, where most cells are actively dividing, providing abundant host replication machinery facilitating efficient viral replication. Previously, it was reported that zika virus preferentially targets proliferating neural progenitor cells, exploiting their high metabolic activity for replication (Gabriel et al., 2017). Furthermore, the innate immune system of zebrafish begins to develop only around day 1 of embryogenesis (Liao et al., 2021; Novoa & Figueras, 2012). The absence of functional innate immune mechanisms in embryos at 0 dpf may allow the introduced hNoV to replicate to high levels before immune sensing becomes effective. Taken together, once the immune system becomes active, a strong antiviral response is mounted, which was observed at 3 dpi in this study. This response was supported by transcriptomic data showing upregulation of pathogen recognition and inflammatory pathways, as well as by metabolomic analyses. The observed perturbation of lipid homeostasis likely reflects hNoV-induced remodelling of membrane composition and lipid-mediated signalling. Such changes may represent a host cell-initiated defense mechanism aimed at limiting viral replication while enhancing immune surveillance (Heaton & Randall, 2011).

The heterogeneity of hNoV infection observed in zebrafish larvae at both the organismal and single-cell levels is highly consistent with previous reports indicating that individual differences in viral infection arise from a dynamic “battle” between viral replication capacity and the host innate immune response (Swaminath and Russell, 2024). At the organism level, genetic background, immune status, and early interferon signalling strongly influence infection outcomes, leading to substantial variation in viral load, disease severity, and clearance kinetics among individuals. At the cellular level, recent single-cell studies reveal that within a seemingly uniform population of cells, viral RNA levels and progeny production can differ by orders of magnitude, reflecting stochastic differences in viral entry, replication efficiency, and host antiviral activation. For example, single-cell transcriptomic analyses of influenza virus infection demonstrated that individual infected cells display highly divergent viral gene expression patterns and innate immune responses, with only a subset of cells activating interferon and ISGs (Sun et al., 2020). Defective or semi-infectious virions can trigger strong innate immune activation in some cells, whereas others support efficient viral replication with minimal antiviral signalling (Russell et al., 2019).

In the present study, scRNA-seq of symptomatic individuals injected with hNoV at 0 dpf revealed viral transcripts across a broad range of cell types at 3 dpi, indicating widespread viral dissemination. Because hNoV was introduced at the early embryonic cell stage, the virus likely had equal opportunity to disseminate into all developing tissues, facilitating systemic spread during development. Furthermore, dextran-AF647 displayed a more diffused distribution in hNoV-infected larvae compared to mock-injected controls, suggesting compromised membrane or barrier integrity. This may reflect virus-induced disruption of epithelial junction and/or inflammatory-related damage, leading to increased tissue permeability (Lechuga & Ivanov, 2018; Troeger et al., 2009). Therefore, widespread viral dissemination likely arises from a combination of early developmental seeding and infection-associated impairment of barrier function. In contrast, only localised infections have been reported in the larval model, with hNoV predominantly concentrated in the intestinal region and caudal hematopoietic tissue (Van Dycke et al, 2019).

Interestingly, scRNA-seq analysis also revealed that majority of virus-containing cells were neuronal cells, suggesting a potential infection of hNoV toward neuronal lineages. Cells associated with the nervous system-associated and those derived from the neural-crest accounted for approximately two-thirds of the total hNoV-infected cell population. Indeed, hNoV infections have been primarily associated with gastroenteritis, but sporadic events linking it to neurological disorder have also been described (Deb et al., 2023). For instance, there have been several reports that linked hNoV infection to gastroenteritis-associated convulsions in children (Chan et al., 2011; Chen et al., 2009; Kawano et al., 2007). In more severe cases, children infected with hNoV developed acute encephalitis (Ito et al., 2006; Obinata et al., 2010; Sanchez-Fauquier et al., 2015). HNoV associated encephalopathy has also been documented in a 60-year-old female patient (Kimura et al., 2010). There have been two reported cases in which hNoV RNA was detected in cerebrospinal fluid of children with encephalitis (Guttierrez-Camus et al., 2022; Ito et al., 2006). While these findings raises the possibility that neural lineages may represent permissive population for hNoV, the mechanisms underlying its entry into and infection of the brain remain poorly understood. On the other hand, our results raise the possibility that hNoV may likely spread to the brain following compromised intestinal epithelial integrity. As observed in zebrafish larvae, a relatively simple organism which systemic infection can occur broadly. In more complex organisms such as humans, dissemination is likely more constrained, with primary infection expected to occur in the intestinal epithelium, followed by barrier disruption that enables viral entry to the bloodstream, subsequent traversal of the blood-brain barrier (BBB), and eventual access into the central nervous system (Ayala-Nunez & Gaudin, 2020). This process may be particularly relevant in vulnerable populations such as children and the elderly, in whom BBB integrity may either be immature or compromised, potentially increasing susceptibility to viral entry into the brain (Bors et al., 2018, Liu et al., 2023; Konig et al., 2025). These findings may have important translational implications for humans, indicating that once hNoV crosses the BBB, it may possess a high potential to cause severe central nervous system infection.

The conventional view holds that non-enveloped viruses, including hNoV, are released only as individual viral particles, principally through cell lysis. However, recent discoveries have shown that non-enveloped virus can also be packaged and non-lytically released from the host cells in the form of EVs (Owusu et al., 2021). EVs have been more and more identified as a mediator for *en bloc* transmission of multiple viruses (Altan-Bonnet et al., 2019; Kerviel et al., 2021). As reviewed by Raab-Traub and Dittmer (2017), EV incorporation can alter the persistence, transmission, and immune recognition of viruses, either promoting or restricting viral replication in target cells. Santiana et al. (2018) reported that EVs were associated with Rotavirus (RoV) and hNoV in stool samples of the hosts, as well as in *in vitro* cell cultivation systems of RoV and MNV as the hNoV surrogate. Although the authors have supplied sound evidence showing that vesicle-contained RoVs from stool are more infectious than free stool rotaviruses both *in vitro* and *in vivo*, it remains unclear whether the same trend applied for hNoVs. In this study, the role of EVs in hNoV infection in zebrafish was demonstrated to be stage dependent. At the early embryonic stage, free viruses generated significantly higher viral loads at 1 dpi and significantly higher symptomatic rates at 3 dpi than EV-associated hNoVs. In addition, GW4869, an inhibitor of EV biogenesis, did not affect the development of pathogenic symptoms in zebrafish, suggesting that free viral particles play a dominant role in driving viral replication and subsequently triggering host immune responses. The relatively lower contribution of vesicle-cloaked viruses may be attributed to factors such as reduced receptor accessibility, altered cellular uptake mechanisms, or delayed uncoating within host cells. Conversely, injection into zebrafish larvae at 3 dpf resulted in the opposite pattern between free and vesicle-cloaked viruses. A subset of individuals injected with free viruses exhibited little to no viral replication, whereas all individuals injected with vesicle-cloaked viruses showed robust replication. We hypothesize that the greater success of vesicle-cloaked hNoV in the more immunocompetent environment of mature larvae may be due to fusion-based uptake into host cells, allowing the virus to evade antiviral immune responses, as reported for several other viruses (Raab-Traub and Dittmer, 2017). As ceramide is also involved in other cellular process beyond EV biogenesis, we therefore examined the effect of inhibiting *de novo* ceramide synthesis using FB1. In contrast to GW4869, FB1 treatment significantly reduced virus-induced pathological symptoms, suggesting that ceramide production promotes hNoV infection independently of EV release. Ceramide is a major component of cellular membranes and plays an important role in regulating membrane organization, fluidity, and the formation of lipid microdomains. These membrane properties are known to influence viral attachment, entry, and intracellular trafficking (Lorizate and Kräusslich, 2011). Moreover, a recent study has demonstrated that hNoV virus-like particle (VLP) clustering and endocytosis are regulated by cholesterol- and ceramide-dependent lipid raft remodeling (Ayyar et al., 2025). Therefore, the reduction in pathological symptoms following FB1 treatment may be attributed to altered membrane architecture that impairs cellular processes required for efficient hNoV infection. In conclusion, the zebrafish embryo model represents a significant advancement in hNoV research by providing a visible proxy for symptomatic infection and disease severity. Our results demonstrate that the timing of infection is critical. The immature immune environment of the 0 dpf embryo allows for systemic viral dissemination and clear pathological manifestations that correlate with viral burden. The identification of hNoV infection for neuronal-related cells via scRNA-seq offers a potential biological basis for the neurological symptoms occasionally observed in human clinical cases. Furthermore, the discovery of stage-dependent roles for EVs suggests that hNoV utilizes diverse transmission strategies to adapt to the host’s immunological maturity. Collectively, this multiscale characterization validates the zebrafish embryo as a highly permissive and versatile model for studying the complex dynamics of hNoV pathogenesis and host-virus interactions.

## Materials and Methods

### Zebrafish maintenance and husbandry

Wild type adult zebrafish (*Danio rerio*) were maintained in the aquatic facility of National University of Singapore with water temperature at 28 °C and 14/10 h light/dark cycle. Fertilized eggs were collected from adults placed in mating cages and kept in petri dishes containing embryo medium E3 (5 mM of NaCl, 0.17 mM KCl, 0.33 mM of CaCl_2_, 0.33 mM MgSO_4_). All zebrafish experiments were performed in compliance with the Institutional Animal Care and Use Committee guidelines, National University of Singapore.

### Microinjection of zebrafish

Fertilised eggs were collected and washed with embryo medium E3 three times. Prior to injection, embryos were inspected under an inverted microscope to ensure that the embryos had not developed past the spherical stage (∼4 hpf). Thereafter, the embryos were transferred to a petri dish with grooves of a mold imprint (6 rows of V-shaped grooves) in 1.5 % agarose. Three nL of the viral suspension was injected into the yolk of each embryo with a pulled borosilicate glass capillary needle. The same steps were performed for larvae injection, except that, 72 h post-fertilised larvae were first anaesthetized for 2 - 3 min in embryo medium E3 containing 0.2 mg/mL Tricaine (Sigma-Aldrich, USA). After which, they were aligned on the agarose surface with their yolk facing the microinjection needle. Three nL of the viral suspension was injected into the yolk of each larva.

### Human norovirus passaging and harvesting

Stool suspension containing hNoV GII.4[P16], kindly provided by the Molecular Laboratory, Department of Molecular Pathology, Singapore General Hospital, was passaged as previously described. Briefly, zebrafish larvae were collected on Day 3 post-injection (3 dpi) and pooled at 10 individuals per sample in a 1.5 mL tube. Larvae were euthanised on ice, the liquid was completely removed, and 100 µL of PBS was added. Thereafter, the pooled larvae were homogenized using by FastPrepTM 24-5G tissue-cell homogenizer (MP Biomedicals, USA) for three cycles of 15 s at 6,500 rpm with rest intervals of 60 s. The resulting homogenates were clarified by centrifugation at 9,000 *×* g for 5 min.

### Embryo collection and dissociation

A total of 20 larvae at 3 dpi were collected and euthanized using tricaine methanesulfonate (MS-222). The larvae were washed twice with phosphate-buffered saline (PBS). Dissociation solution comprising 480 µL of 0.25% trypsin and 20 µL of 100 mg/mL collagenase (total volume: 500 µL) was added to the washed larvae. Mechanical dissociation was performed by pipetting the mixture for 30 s, followed by incubation at 30 °C for 30 s. This cycle was repeated until the tissue was fully dissociated and no visible fragments remained, typically within 5 min. Enzymatic digestion was terminated by adding 800 µL of Dulbecco’s Modified Eagle Medium supplemented with 10% fetal bovine serum (DMEM–10% FBS). The cell suspension was then centrifuged at 700 *× g* for 5 min at room temperature. The supernatant was discarded, and the pellet was washed once with PBS. The washed cell pellet was resuspended in 500 µL of PBS with 0.04% bovine serum albumin (BSA) and passed through a 40 µm cell strainer to obtain a single-cell suspension.

### RNA extraction and RT-qPCR for hNoV detection

RNA was extracted using RNeasy Mini Kit (Qiagen, Germany) following the manufacturer’s protocol. RT-qPCR analyses of NoV GII were carried out using GoTaq® Probe 1-Step RT-qPCR System (Promega, USA). Primers and the FAM/TAMRA-labelled probe of hNoV GII were according to ISO 15216-1:2017. Forward primer QNIF2: 5’-ATGTTCAGRTGGATGAGRTTCTCWGA-3’; reverse primer G2SKR: 5’-TCGACGCCATCTTCATTCACA-3’; probe QNIFs: 5’ FAM-AGCACGTGGGAGGGCGATCG-3’ TAMRA. Cycling conditions were 45°C for 15 min, then 95°C for 10 min, followed by 40 cycles with 95 °C for 15 s and 60 °C for 30 s in each cycle. Cycle threshold (Ct) values were determined during RT-qPCR analysis using StepOneTM system (Applied Biosystems, USA).

Double-stranded DNA (dsDNA) containing the specific primers-probe binding sites were synthesized for NoV GII and cloned into the pGEM-T Vector (Promega) resulting in the NoV-GII plasmids. The plasmids with inserts were purified by using a Plasmid Midi Kit (Qiagen). The plasmid concentration was determined by photospectroscopy at 260 nm using the BioDrop DuoTM spectrophotometer (BioDrop, United Kingdom). Ten-fold serial dilutions ranging from 5 × 10^6^ to 5 copies of all positive control plasmids were used to prepare a standard curve and enumerate the NoV GII.

### RNA sequencing and sequencing analysis

mRNA was purified from total RNA using poly-T oligo-attached magnetic beads. After fragmentation, the first strand cDNA was synthesized using random hexamer primers. Then the second strand cDNA was synthesized using dUTP. The directional library was ready after end repair, A-tailing, adapter ligation, size selection, USER enzyme digestion, amplification, and purification. The library was checked with Qubit and real-time PCR for quantification and bioanalyzer for size distribution detection. Quantified libraries were pooled and sequenced on NovaSeq6000 (Illumina) PE150 using paired-end 150 run (2 × 150), according to effective library concentration and data amount.

Sequencing data were analysed using the Galaxy web platform (https://usegalaxy.org/) (Afgan et al., 2016). Raw reads were screened with FastQC, followed by trimming with Trimmomatic to remove adapters and low-quality reads (Bolger et al., 2014). Trimmed reads were aligned with Spliced Transcripts Alignment to a Reference (STAR) to zebrafish (Danio rerio) genome build, GRCz11; retrieved from Ensembl. The gene-level read counts were measured with featureCounts (Liao et al., 2013). Differential gene expression was analyzed with DESeq2 (Love et al., 2014), and functional enrichment analysis was conducted using ShinyGO v.0.85.

### Single-cell library preparation and sequencing

Library preparation was performed using Chromium™ Next GEM Single Cell 3’ GEM, Library & Gel Bead Kit v3.1. Single cell 3’ RNAseq experiment was conducted using the Chromium single cell system (10x Genomics) and the NovaSeq 6000 sequencer (Illumina). Following cell capture and lysis, cDNA was synthesized and amplified, and libraries were subsequently prepared from the amplified cDNA. The library was checked with Qubit and real-time PCR for quantification and bioanalyzer for size distribution detection. Quantified libraries were pooled and sequenced on NovaSeq6000 PE150 using paired-end 150 run (2 × 150), according to effective library concentration and data amount.

### Single-cell data processing

Reads were analyzed using Cell Ranger (10x Genomics). STAR was used to align the reads to zebrafish (Danio rerio) genome build, GRCz11. Quality control and downstream analysis were performed using Seurat. Low quality cells were excluded with the following criteria: cells with gene counts less than 200 or over 5000, >50% mitochondrial genes, or >5% Heparin-Binding genes. Subsequently, DoubletFinder was used to identify and remove doublets, and only singlets were retained for downstream analysis.

Cells were clustered using the principal component analysis (PCA) and visualised using the uniform manifold approximation and projection (UMAP) plot. FindVariableGenes function was used to identify highly variable genes, and the top 3000 most variable genes from each sample were selected as input for dimensional reduction via PCA with RunPCA function. The top 15 principal components, selected based on the elbow point, were then used for cell clustering with FindCluster function. The differentially expressed genes (DEGs) between each cluster were calculated with the FindAllMarker function using Wilcoxon rank sum test with the threshold q-value < 0.05 and log2FC > 0.25. Cells were annotated using the FindAllMarkers function, referencing the expression profiles of the top 16 most differentially expressed genes previously characterized by Farnsworth et al. (2019).

### Metabolite extraction from larvae

Ten larvae at 3 dpi were pooled to form one biological replicate, then euthanised and washed as previously described. Each group was immediately homogenized in 1 mL of ice-cold LC-MS grade methanol (Waters, Milford, MA, USA). To achieve complete cell lysis and metabolite release, the homogenate was subjected to 3 freeze-thaw cycles using liquid nitrogen and thawing on ice. The samples were then incubated at −20 °C overnight to facilitate protein precipitation. The following day, the samples were centrifuged at 12,000 *× g* for 20 min at 4 °C. The resulting supernatants were collected and mixed with 0.1 mM gallic acid (as an internal standard). The mixture was then filtered through a 0.22 μm nylon membrane filter (Waters, Milford, MA, USA) and transferred to LC-MS vials for analysis. A pooled quality control (QC) sample was prepared by combining equal aliquots from each experimental group and was injected periodically throughout the run to monitor instrument stability and analytical reproducibility.

### UPLC-QTOF-MS Analysis

Metabolomic profiling was performed using an ACQUITY UPLC™ I-Class PLUS system (Waters, Milford, MA, USA) coupled with a Xevo G2-XS quadrupole time-of-flight mass spectrometer (QTOF-MS; Waters, Manchester, UK). The system was equipped with an electrospray ionization (ESI) source operated in both positive (ESI+) and negative (ESI−) ion modes. Chromatographic separation was achieved using an ACQUITY UPLC BEH C18 column (2.1 × 100 mm, 1.7 μm; Waters) maintained at 40 °C. The mobile phase consisted of 0.1% formic acid in water (solvent A) and 0.1% formic acid in acetonitrile (solvent B). The elution was performed with the following gradient program: 0–1 min, 5% B; 1–12 min, linear increase to 95% B; 12–15 min, held at 95% B; 15–15.5 min, returned to 5% B; and 15.5–18 min, re-equilibrated at 5% B. The flow rate was set to 0.3 mL/min, and the injection volume was 5 μL. The QTOF-MS conditions were as follows: capillary voltage, 2.5 kV; source temperature, 120 °C; desolvation temperature, 450 °C; desolvation gas flow, 800 L/h. Data were acquired in centroid mode over a mass range of m/z 50–1200. Leucine-enkephalin was used as the reference lock mass to ensure mass accuracy throughout the run.

### Spectral analysis

The Progenesis QI program (V.2.4, Nonlinear Dynamics, Waters, Newcastle, U.K.) was used to analyze the UPLC-QTOF-MS spectra, which automatically carried out retention time alignment, baseline filtering, response normalization, and 3D peak selection. The adducts [M + H]+, [M + NH4]+, [M + Na]+, and [M + K]+ were chosen for the positive mode of metabolite identification, while [M - H]- and [M + FA - H]- were chosen for the negative mode. The Kyoto Encyclopedia of Genes and Genomes (KEGG), and the Human Metabolome Database (HMDB), were all used as references for metabolite identification. Further multivariate analysis was performed using orthogonal partial least squares discriminant analysis (OPLS-DA) with EZinfo software (V.3.0, Umetrics, Sweden). Substantially changed metabolites in the treatment group compared with the control group (VIP > 1, *P* < 0.05, and coefficient of variation [CV] ≤ 30%) were considered as significant. To visualize the differential metabolite profiles, a hierarchical clustering heatmap was generated using TBtools software (v1.106). Prior to visualization, the data matrix of significant metabolites was log2-transformed and normalized by Z-score scaling. Clustering was performed based on Euclidean distance and average linkage method. Both metabolite-wise and sample-wise dendrograms were constructed to reveal group-specific metabolic patterns.

### Extracellular vesicle isolation

Extracellular vesicles were isolated from clarified virus suspension using MagCapture Exosome Isolation Kit PS using TIM4 coated magnetic beads (Fujifilm, Japan) according to manufacturer’s protocol. Briefly, samples were incubated with TIM4 coated magnetic beads for an hour at RT with gentle mixing to allow vesicles to bind to the beads. The beads-vesicles complex was then separated from the sample, using a magnetic trip for 1 min at RT, and washed 3 times with the buffer included in the kit. Finally, the bead bound vesicles were eluted from the beads for further processing.

### Virus purification by sucrose cushion low-speed centrifugation

Clarified virus suspension was further purified by centrifugation through a 20 % (w/v) sucrose cushion. Briefly, virus suspension was carefully layered onto a 20 % sucrose cushion prepared in PBS. Samples were then centrifuged at 18,000 *× g* for 20 h at 4 °C using a Thermo Fisher benchtop centrifuge with 1.5 mL Eppendorf tubes. After centrifugation, the supernatant was discarded, and the viral pellet was resuspended in PBS. To concentrate the virus, the resuspended sample was filtered through a 100 kDa molecular weight cut-off (MWCO) centrifugal filter unit (Millipore). The concentrated virus was stored at 4 °C until further use.

### Microscopic imaging

Infected larvae were anaesthetized for 2 - 3 min in embryo medium E3 containing 0.2 mg/mL Tricaine (Sigma-Aldrich, USA). After which, they were embedded in 3% methylcellulose to immobilise them for imaging. Images were captured using a Nikon SMZ25 stereo microscope (Nikon, Japan) equipped with Nikon DS-10 digital camera.

For the membrane integrity assay, embryos were co-injected with 1mg/ml of 10 kda Dextran, Alexa Fluor™ 647 (Thermo Fisher Scientific, USA). Larvae were visualized using the TexasRed Channel (AT560/40x excitation filter, AT635/60m emission filter, AT600DC dichroic mirror) with a 2 s exposure time for fluorescence image acquisition

### Cryo-Electron Microscopy (Cryo-EM)

Three µL of concentrated and filtered larva lysis was applied to glow-discharged cryo-EM grids (CF-2/1-3CU, Protochips). The grids were plunge-frozen in liquid ethane using a Leica EM GP or Leica EM GP2 and stored in liquid nitrogen for subsequent imaging. Cryo-EM grids were loaded into a Titan Krios transmission electron microscope (TEM) (Thermo Fisher Scientific) operated at 300 kV and equipped with a BioContinuum K3 direct electron detector. Images were recorded using SerialEM-4.0.6.

### Ceramide pathway inhibition treatment

Viral-injected embryos were treated with GW4869 (Sigma-Aldrich, USA) and Fumonisin B1 (FB1; Sigma-Aldrich, USA), respectively, to inhibit different pathways involved in ceramide production. Toxicity ranges for both compounds were initially screened, and the highest non-toxic concentrations were selected for treatment. The concentration used were 20µM for GW4869 and 5µM for FB1. Treatments were maintained in E3 medium for a duration of 3 days.

For easier visualization under the microscope, larvae were individually transferred into the wells of a 96-well plate at 2 dpi. Subsequently, at 3dpi, the individual larvae were anaesthetized for 2 - 3 min in embryo medium E3 containing 0.2 mg/mL Tricaine (Sigma-Aldrich, USA) to facilitate observation of the pathological symptoms.

### Statistical analysis

Statistical analyses were performed using the software SPSS for windows, version 31.0. Student’s t-test was used to compare the means between two independent groups, and one-way analysis of variance (ANOVA) was used for data with more than two independent groups, the Tukey test was used as post-hoc test. Significant differences were considered when *P* value < 0.05.

## Acknowledgements

This study made use of electron microscopes at the NUS Centre for Bioimaging Sciences (CBIS) and we thank Jian Shi for technical assistance. This study was financially supported by Department of Food Science and Technology, National University of Singapore (PI: Dan Li, E-160-00-0015-001), and the National University of Singapore Presidential Young Professorship start-up grant (PI: Kun Qu, NUHSRO/2022/033/Startup/03).

## Author Contributions

**M.T.H.T.** and **D.L.** conceived and designed the study. **M.T.H.T.**, together with **H.D., Z.L., J.Y.L.T.,** and **H.B**., performed the experiments. **M.T.H.T.** analyzed the data, and wrote the original draft. **D.L.** and **Q.K.** provided resources, supervised the project, and critically revised the manuscript. All authors read and approved the final manuscript.

